# Snapshots of single particles from single cells using electron microscopy

**DOI:** 10.1101/435222

**Authors:** Xiunan Yi, Eric J. Verbeke, Yiran Chang, Daniel J. Dickinson, David W. Taylor

## Abstract

Cryo-electron microscopy has become an indispensable tool for structural studies of biological macromolecules. There are two predominant methods for studying the architectures of multi-protein complexes: (1) single particle analysis of purified samples and (2) tomography of whole cells or cell sections. The former can produce high-resolution structures but is limited to highly purified samples, while the latter can capture proteins in their native state but is hindered by a low signal-to-noise ratio and results in lower-resolution structures. Here, we present a method combining microfluidic single cell extraction with single particle analysis by electron microscopy to characterize protein complexes from individual *C. elegans* embryos. Using this approach, we uncover three-dimensional structures of ribosomes directly from single embryo extracts. In addition, we investigate structural dynamics during development by counting the number of ribosomes per polysome in early and late embyros. This approach has significant potential applications for counting protein complexes and studying protein architectures from single cells in developmental, evolutionary and disease contexts.

## Introduction

Cell behavior is fundamentally dependent on the activities of macromolecular machines. These machines, comprised of protein (and sometimes RNA) subunits, are responsible for catalytic, structural and regulatory activities that allow cells to function. Structural biology, by revealing the physical architecture of macromolecules and their assemblies, plays a critical role in efforts to understand how molecular mechanisms contribute to cell behavior *in vivo*.

A crucial feature of most living cells is their ability to adjust their behavior in response to their environment. In a developmental context, cells respond to chemical and mechanical cues from neighboring cells and tissues in order to coordinate their behavior with their neighbors and to assemble functional tissues. A major goal of developmental biology studies is to understand the molecular mechanisms of these interactions – that is, how dynamic behaviors of macromolecular machines give rise to cell behaviors that support proper organismal development.

Recently, single cell nucleic acid sequencing approaches have revolutionized developmental studies by allowing gene expression to be interrogated with unprecedented spatiotemporal resolution (Kumar et al., 2017). Such experiments are powerful because they reveal which genes are expressed in which cells at a particular point in development, and can thus provide insights into signaling dynamics, mechanisms of cell state changes (e.g. cellular differentiation) and levels of heterogeneity between individual cells. However, sequencing approaches do not shed light on the molecular states or interactions of cellular proteins. A few studies have begun to extend a single-cell approach to biochemical studies of proteins and protein complexes. For example, Huang and Zare (2007) described a sophisticated microfluidic device for counting protein molecules in single-cell lysates. Allbritton and colleagues have developed capillary electrophoreses methods for measuring enzyme activities in whole-cell lysates (Kovarik, 2011; Dickinson et al., 2013). Most recently, Dickinson and colleagues (2017) used microfluidic lysis followed by single-molecule pull down and TIRF microscopy to measure the abundance of protein complexes in single cells. This <u>s</u>ingle-<u>c</u>ell, <u>s</u>ingle-<u>m</u>olecule <u>pull</u>-down (sc-SiMPull) approach was sufficiently sensitive to reveal regulated changes in protein-protein interactions that occurred over ~5 minutes during development of the *Caenorhabditis elegans* zygote (Dickinson et al., 2017). Thus, single-cell biochemical approaches have the potential to uncover dynamics of macromolecular machines in cell or tissue samples obtained directly from developing embryos.

Although still in their infancy, the initial success of these single-cell biochemical methods raises the question of whether a single-cell approach could be extended to macromolecular structure determination. Such an approach could overcome a classical limitation of structural biology: its need for highly purified, homogenous proteins (or protein complexes) that represent only a single snapshot from the ensemble of structures that are likely present in cells. Moreover, the ability to determine structures of proteins obtained directly from cells engaged in development would represent a significant step towards the goal of linking the structural dynamics of molecular machines to their cellular and developmental consequences.

One approach to single cell structural studies is electron tomography. This allows for the study of cell morphologies (Beck and Baumeister, 2016), and in some cases, can be used to reconstruct three-dimensional (3D) models directly from native cells (Galaz-Montoya and Ludtke, 2017). However, due to the sensitivity of biological specimens to electron dose, tomographic approaches routinely lead to low-or intermediate-resolution structures of complexes in single cells using sub-tomogram averaging techniques. The advent of phase plates for electron microscopy has revolutionized the information content extractable from tomograms, but high-resolution structures of less-abundant complexes remain elusive.

Alternatively, single particle cryo-electron microscopy (cryo-EM) is now capable of routinely achieving high-resolution structures of highly purified samples due to advances in hardware (Kühlbrandt, 2014) and software (Scheres, 2012; Punjani et al., 2017). We and others have recently extended single particle EM techniques to study heterogenous mixtures from biochemically fractionated cell lysate (Kastritis et al., 2017; Verbeke et al., 2018). While these shotgun-EM approaches are able to sort through the heterogeneity of macromolecules, they still rely on a large quantity of cells and mass spectrometry to characterize the contents of the sample. Furthermore, direct investigation of proteins at the single cell level has remained a challenging problem for proteomic studies. This poses a unique challenge to structural studies of single cells.

To reconcile single cell and single particle methods, we propose an alternative approach which combines single cell lysis with EM to investigate individual *C. elegans* embryos. Using this method, we are able to directly visualize the contents of a single *C. elegans* cell (the zygote). After computationally classifying the particles from cell lysate, we uncover the structures of 40S and 60S ribosomes from disperse particles and the structure of an 80S ribosome from polysomes. This approach additionally enables us to count individual particles from embryos at specific developmental stages. In one application, we find that the number of ribosomes per polysome remains consistent between early and late stage embryos. These results demonstrate the potential of EM for structural characterization of unpurified macromolecular machines obtained from samples as small as a single cell.

## Results

### Extracting macromolecules from single embryos

Our primary goal in this study was to determine whether imaging of single cell lysates with EM could yield sufficient, high-quality particles for 3D structure determination. To obtain intact, native particles from single cells, *C. elegans* zygotes (i.e., 1-cell embryos) were trapped and lysed using microfluidic chambers (Figure 1) (see Methods). We then transferred the cell lysates (a volume of ~50 nL) from the microfluidic channels to EM grids using a glass needle (Supplemental Movie 1) (see Methods). Due to the small volume, which was insufficient to coat an entire grid, we used reference grids containing alphanumeric markers to locate the placement of our samples under both the dissecting scope and the electron microscope. Each reference grid was then conventionally stained using 2% (w/v) uranyl acetate. We chose to use negative stain EM for its high-signal-to-noise ratio in order to more accurately assess our ability to identify single particles from individual cell lysates. Each grid was then examined by transmission EM to identify grid squares that contained cellular protein particles embedded in stain.

**Figure 1.**
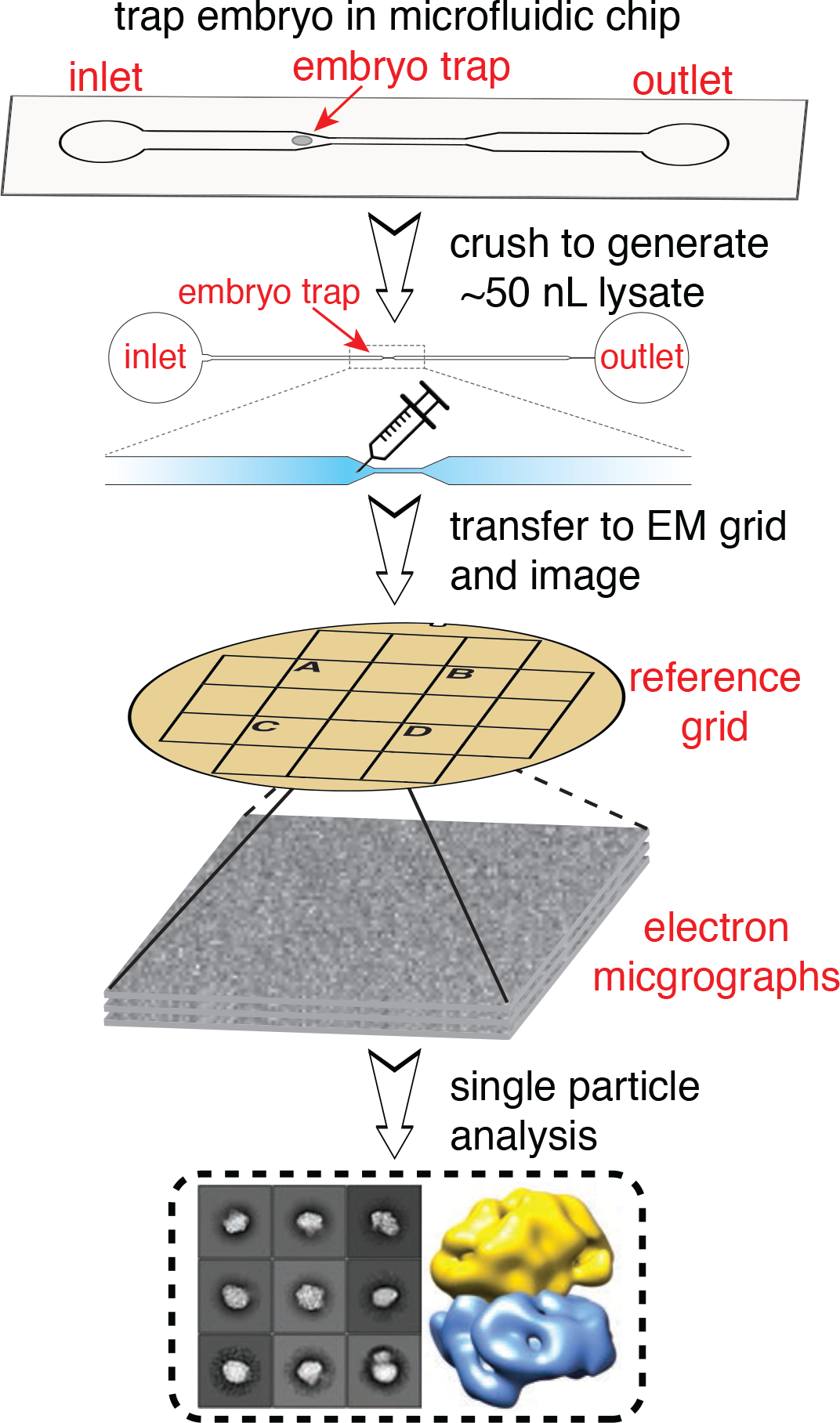
Schematic of single cell structural biology approach. Single *C. elegans* embryos are trapped in a microfluidic device. After the embryo is crushed, the lysate is extracted using a fine needle and applied to a specific area of a EM grid using a stereoscope. The same area is then visualized using electron microscopy and single particle analysis is applied for structure determination.

To demonstrate our ability to capture small volumes of samples on EM grids, we first transferred samples of a purified protein kinase from our microfluidic device to an EM grid for visualization, resulting in successful detection of the kinase (Figure S1). We then performed our transfer technique on lysates from seven, independent single embryos sampled at different developmental stages, 3 from early-stage and 4 from late-stage. Micrographs of single cell extract across different embryos show a reproducible mixture of heterogeneous particles which span an order of magnitude in size (Figure S1). This data allowed us to investigate the dynamics of protein complexes at the single cell/embryo level between different developmental stages of *C. elegans* zygotes.

### Electron microscopy of extract from a single*C. elegans* embryo

Raw micrographs from data collected at the locations where embryo lysate had been applied to the EM grid showed distinct, monodisperse particles with varying sizes and distinct shapes. The results unambiguously show that we were able to retrieve cellular contents from our microfluidic lysis chips for subsequent imaging by EM (Figure 2A). We collected ~1,400 micrographs between the 7 samples. While small particles were abundant in our micrographs (Figure S1), we first chose to analyze large particles (~150-300 Å in diameter), which were easily recognizable and appeared relatively homogeneous. After manually selecting ~10,000 large particles, we generated reference-free 2D class averages, which were subsequently used as templates for picking the remainder of the particles. Using this template picking scheme, ~80,000 large particles were selected from ~1,400 micrographs and used for reference-free 2D alignment and classification. 2D class averages with distinct structural features were generated from ~50,000 particles after removing junk particles (e.g. detergent micelles, irregular small particles, two nearby particles, or particles in aggregates) from the data (Figure 2B Top).

**Figure 2.**
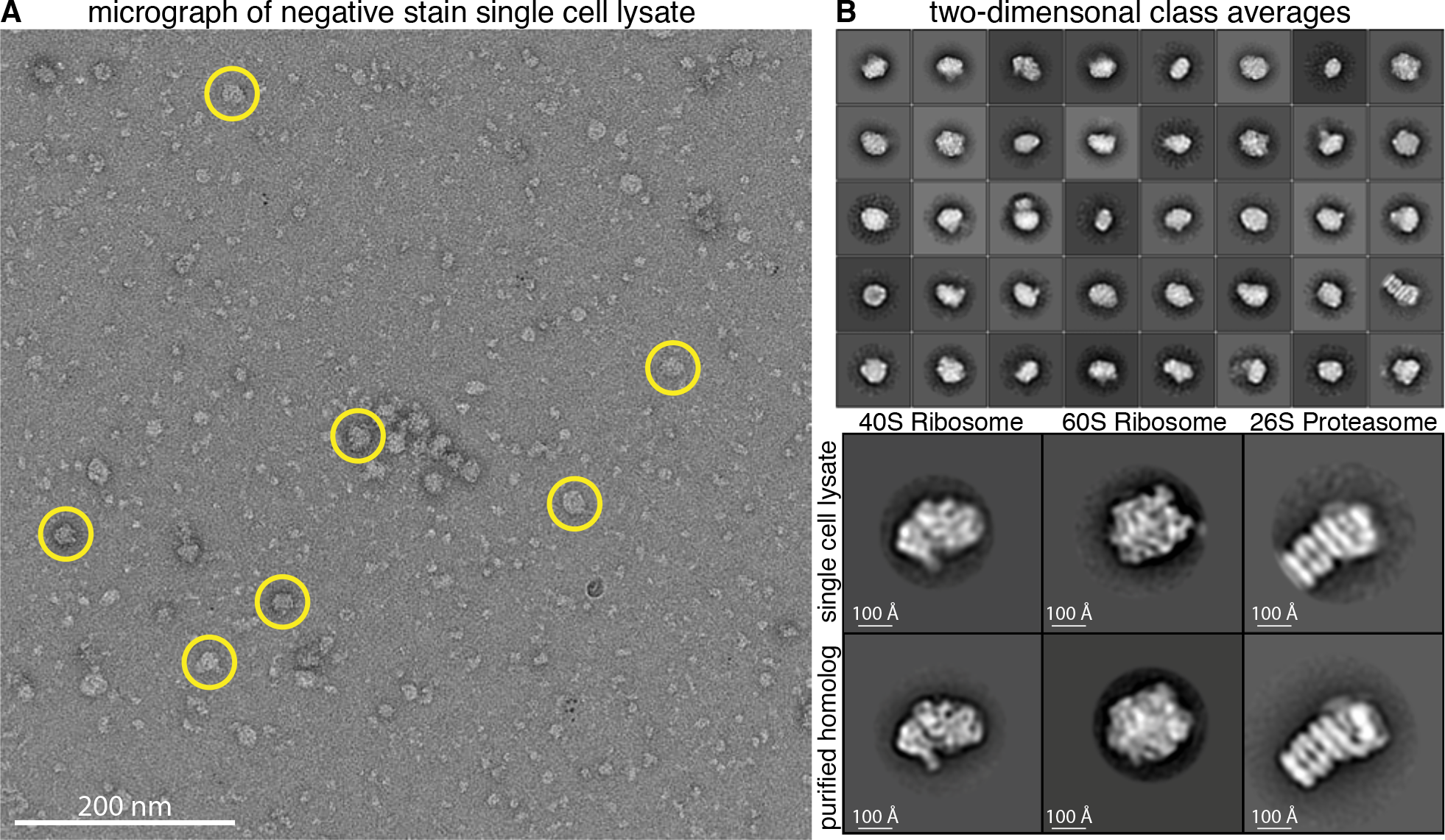
Single particle analysis of extracts from single cells. Representative raw electron micrograph of negatively stained single cell lysates. Micrographs show monodisperse particles of varying size. Circled particles are representative of the larger particles (~150-300 Å in diameter) used for subsequent two-and three-dimensional classification.(B)Top: Reference-free 2D alignment and classification of a subset of the ~50,000 particles picked from single cell extract. Classes are sorted in order of decreasing abundance. Box size is 576 Å x 576 Å. Bottom: Alignment of 2D class averages from single cell extract to purified homologs.

To obtain insight into the possible identities of these particles, we used publicly-available RNA-seq data from *C. elegans* 1-cell embryos (Hillier et al., 2009; Gerstein et al., 2010) to inform us about which proteins are likely to be highly expressed. 34 ribosomal protein transcripts and 4 proteasomal protein transcripts were among the 200 most abundant transcripts, suggesting these protein complexes were likely to appear in our micrographs (Figure S2) (Hillier et al., 2009; Gerstein et al., 2010). We therefore performed pairwise cross-correlations of our 2D class averages with 2D class averages of purified 40S ribosome, 60S ribosome, and 26S proteasome from *S. cerevisiae* to look for structural similarities. The alignment revealed several classes with similar features between our single-cell lysate and the known, purified structures, suggesting the identity of several projections in our sample were in fact the 40S ribosome, 60S ribosome, and 26S proteasome (Figure 2B Bottom). This initial 2D classification proved it is possible to obtain structural information from intact protein complexes extracted from lysates of single cells.

### Capturing ribosome dynamics in polysomes

Intriguingly, our raw micrographs revealed densely-packed clusters of ribosome-like particles (Figure 3A). These ribosome-like particles appeared in organized arrays with a similar appearance to polysomes from *E. coli* (Brandt et al., 2009) and wheat germ (Afonina et al., 2013). Polysomes consist of a pool of actively translating ribosomes on an mRNA transcript. This suggested that our single-cell EM method is capable of capturing protein-mRNA interactions in the cell.

**Figure 3:**
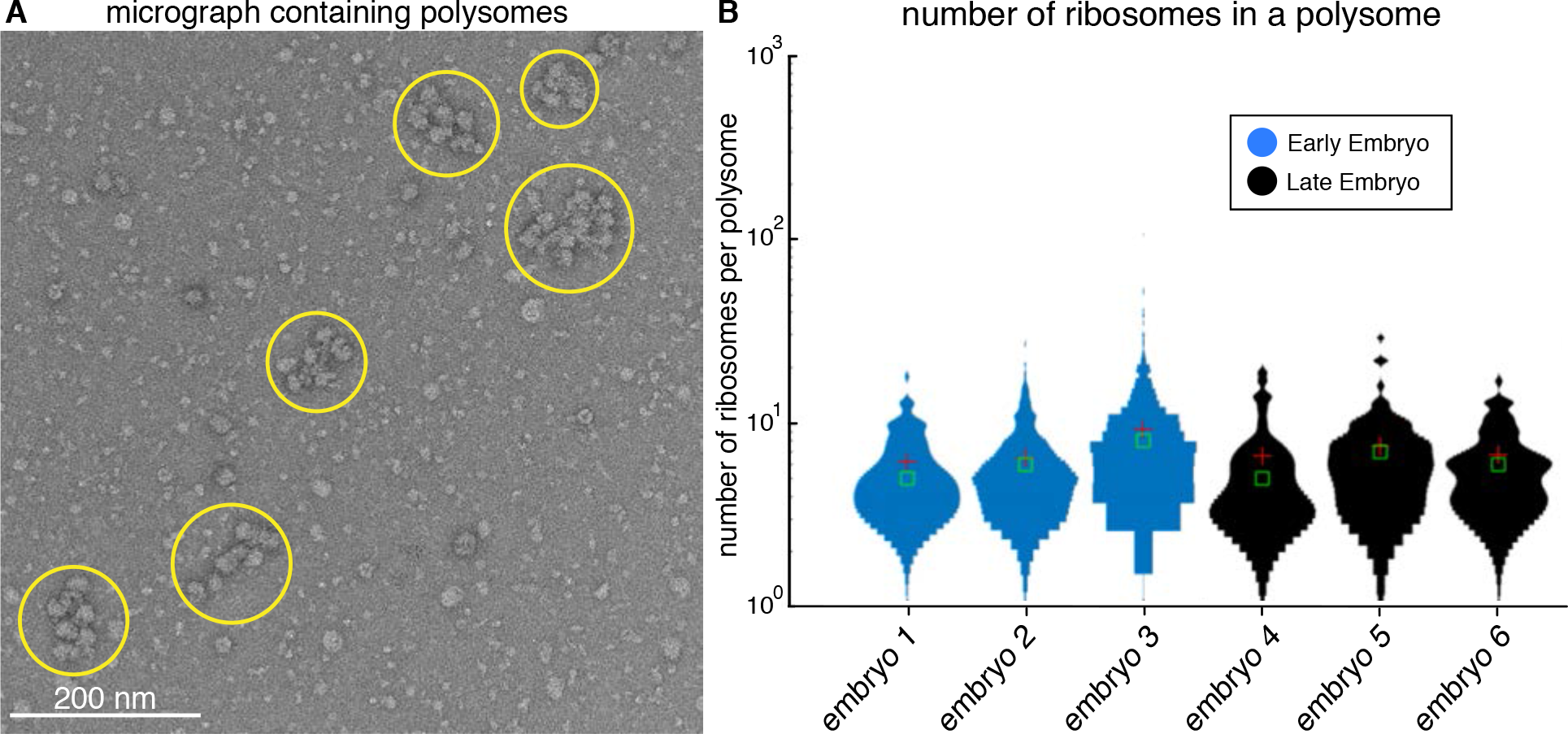
Counting ribosomes in polysomes from early-and late-stage*C. elegans* embryos. (A)Representative raw electron micrograph of negatively stained single cell lysate showing several distinct polysome clusters of varying size (yellow circles). (B)Distribution of the number of ribosomes in a polysome across 3 early-and 3 late-stage embryos. The average number of ribosomes for early-and late-stage embryos is 8 and 7, respectively. The red crosshair is the mean value and the green box is the median.

We were interested in observing polysome dynamics through developmental stages of embryos by counting the number of ribosomes per polysome. Each micrograph was manually annotated to determine the number of ribosomes per polysome cluster for our early-stage and late-stage embryos, respectively. Using this approach, we determined there are on average 8 ribosomes per polysome for early-stage embryos and 7 ribosomes per polysome for late-stage embryos (Figure 3B). These numbers are consistent with previous studies in which the number of ribosome per mRNA is estimated by isolating polysomes using velocity sedimentation in sucrose gradients (Slavov et al., 2015). Thus, our data suggested no significant change in the number of ribosome per polysome between early and late stage embryos. This analysis provides evidence that single particle counting from single cells could potentially be applied for investigating the dynamics of macromolecules in different cell states.

### 3D classification of ribosome particles from single embryo data

We then performed 3D classification of our large particles to determine if any distinct structures could be obtained from lysates of single cells. Specifically, we were looking for structures of ribosomes since they appeared as clear and abundant 2D class averages in our data. We first combined two datasets from early-stage embryo samples for 3D classification using RELION (Scheres, 2012). After removal of junk particles, ~14,000 particles were used for classification. Initially, we used an unbiased approach for 3D classification by using an initial model of a featureless 3D shape with uniform electron density. Using a model reconstructed from this initial classification that resembled a previously determined 60S ribosome structure (Shen et al., 2015) (EMDB-2811) as a reference, we then performed another round of 3D classification (see Methods). The models from each classification were then compared by docking a high-resolution *S. cerevisiae* 60S ribosome structure (EMDB-2811) into our maps to determine which class, if any, was most similar to the known structure. Our top scoring 60S ribosome reconstruction, containing ~3,400 particles, displayed striking similarity to the *S. cerevisiae* 60S ribosome (Figure 4 Top) with a cross-correlation score of 0.8143 and a nominal resolution of 34 Å calculated using the 0.5 Fourier shell correlation criterion (Figure S3) (see Methods).

**Figure 4:**
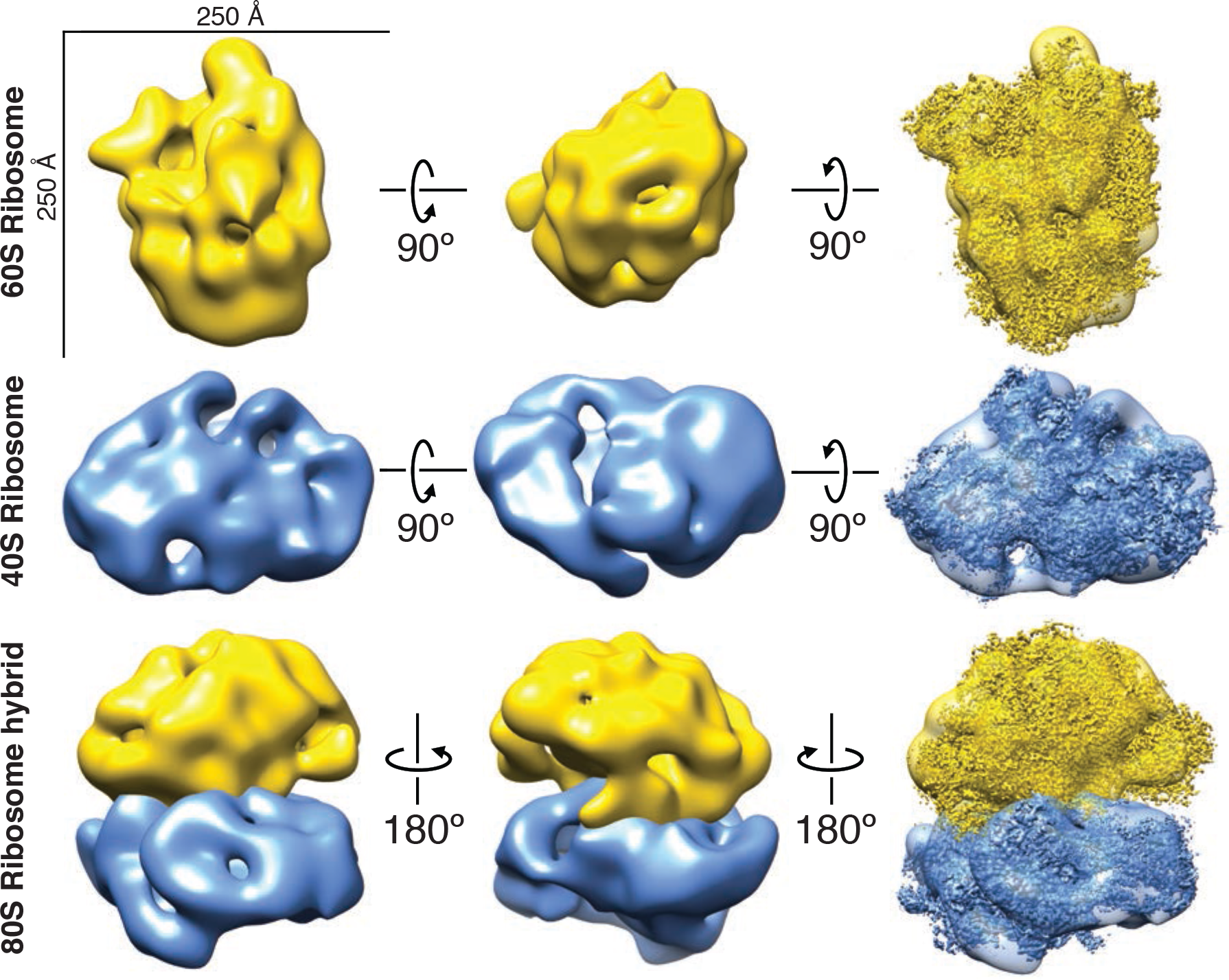
40S and 60S ribosome reconstructions from particles from single cells. Top: 60S ribosome reconstruction. High-resolution structure EMDB-2811 (Shen *et al.*, 2015) docked into our 60S map with a cross-correlation score of 0.8142. Middle: 40S ribosome reconstruction. High-resolution structure EMDB-4214 (Scaiola *et al.*, 2018) docked in to our 40S map with a cross-correlation score of 0.8352. Bottom: 80S ribosome hybrid model built using our 40S and 60S ribosome aligned to a high-resolution structure of the 80S ribosome EMDB-2858 (Cianfrocco and Leschziner, 2015).

Performing 3D classification on particles from all datasets combined, a final set of ~17,000 particles after stringent removal of junk particles, resulted in an additional class which resembled the *S. cerevisiae* 40S ribosome (Figure 4 Middle) (see Methods). Our 40S ribosome reconstruction, containing ~1,450 particles, had a cross-correlation score of 0.8352 with the *S. cerevisiae* model (Scaiola et al., 2018) (EMDB-4214) and a nominal resolution of 48 Å (Figure S3). These results suggest that 3D structures of multiple protein complexes can be obtained from lysates of single embryos using single-particle EM analysis. To explore our ribosome reconstructions, we built a hybrid 80S ribosome model by aligning our 40S and 60S ribosome reconstructions to their respective domains in the 80S ribosome from a previously determined 80S ribosome structure (Figure 4 Bottom) (Cianfrocco and Leschziner, 2015) (EMDB-2858). As expected, this hybrid model is consistent with the high-resolution 80S ribosome structure.

With clear structures resembling a 40S and 60S ribosome, we next attempted to determine the molecular architecture of the 80S ribosome directly from polysome clusters. Our data contained ~9,000 particles within polysomes which were manually picked for single particle analysis. We then performed 2D and 3D classification of the selected particles (see Methods). The 2D class averages were approximately 250 Å in diameter, which is consistent with the size of an *S. cerevisiae* 80S ribosome. Our 3D model, containing ~2,000 particles, had a cross correlation score of 0.7572 when compared to an *S. cerevisiae* 80S ribosome (Cianfrocco and Leschziner, 2015) (EMDB-2858) and a resolution of 45 Å (Figure S3). Although our 80S ribosome model lacked some areas of density present in the high-resolution structure, the overall size could accommodate both the 40S and 60S ribosome. Collectively our data shows that we are able to distinguish structures of the 40S, 60S and 80S ribosome directly from particles isolated from single cells.

## Discussion

A major goal of basic biological research is to connect structural dynamics of macromolecules to their effects on cell behavior. Here, we present an approach for structural characterization of protein complexes isolated from single cells engaged in development. We demonstrate that a single cell contains a sufficient number of protein particles to enable structural characterization by EM. We think that this approach has significant potential to reveal structural changes in protein complexes across developmental and disease contexts.

Moving forward, a significant challenge will be to extend this approach beyond ribosomes, to other macromolecular complexes. We focused here on ribosomes because they are large, highly abundant, and relatively easy to recognize. For complexes that are less abundant and/or less distinctive in shape, we will need to develop methods to identify a complex of interest in a heterogenous mixture. Correlative Light and Electron Microscopy (CLEM) holds promise in this regard (Schorb and Briggs, 2014). We also plan to explore whether particles isolated and characterized via our earlier sc-SiMPull approach (Dickinson et al., 2017) can be eluted and transferred to EM grids for structural analysis. An added advantage of this strategy would be the ability to use multicolor TIRF to characterize the composition of complexes whose structures could then be determined. Taken together, we are optimistic that these strategies will allow us to gain structural information about any protein complex of interest using single-cell lysates.

A related question is whether there are enough particles in a single cell to allow high-resolution structure determination. This will of course depend on the protein or protein complex being studied; it may be more difficult to determine high-resolution structures of low-abundance complexes. However, we note that the time required to prepare and collect data from a single-cell sample is short enough that analyzing 10-20 such samples is realistic. Particles from multiple samples could be pooled to increase resolution, without sacrificing information about which cell each particle in the dataset came from. This might represent an ideal compromise between the need for increased numbers of particles for structure determination and the desire for single-cell resolution for detailed developmental studies.

## Author Contributions

X.Y. and Y.C. performed sample preparation of embryo lysate. X.Y., E.J.V., and Y.C. performed electron microscopy, single particle classification and reconstruction. X.Y., E.J.V., Y.C., D.J.D., and D.W.T. analyzed and interpreted the data and wrote the manuscript. D.J.D. and D.W.T. conceived of experiments. D.J.D. and D.W.T. supervised the study and secured funding for the work.

## Methods

### Microfluidic device fabrication

Microfluidic devices were fabricated using a standard soft lithography procedure. A photomask corresponding to the desired channel shape was designed using CAD software and produced by Cad-Art Services (Bandon, OR, USA). An approximately 30 *μ*m-thick layer of SU8-2025 photoresist was deposited on a plasma-treated silicon wafer by spin coating for 10s at 400 rpm followed by 30s at 2800 rpm and 30s of deceleration. After soft baking at 65° for 3min and 95° for 10 min., the films were exposed to 1000 mJ UV light through the photomask. Following a post-exposure bake of 5 min. each at 95° and 120°, the molds were developed in SU8 developer (propylene glycol monomethyl ether acetate, PGMEA) and rinsed with isopropanol. The molds were hard baked at 95° for 30 minutes and then at 120° overnight.

PDMS (Sylgard 184 silicone elastomer kit, Dow Corning, Midland, MI) was mixed using a 10:1 ratio of base to curing agent and deposited onto the molds by spin coating at 400 rpm for 30s. The PDMS was cured for 20 minutes at 95°, then peeled off from the molds, and inlet and outlet holes were punched with a 2 mm biopsy punch. Each PDMS device contained 8 channels, and each channel was used for one single-embryo experiment.

24×60mm glass coverslips were cleaned with ethanol and dried under nitrogen flow. Each cleaned coverslip was bonded to a PDMS device by 2 min. treatment with air plasma, then baked at 120° for 30 minutes to form a permanent bond.

The PDMS device was first activated by flowing 1 M KOH through the channels for 20 min., washed 3 times with water, and then dried. After activation, 2-[methoxy(polyethylenxy)9-12Propyl]-trimethoxysilane was applied to the channels for 30min. to prevent non-specific protein binding. The channels were then washed 3 times with water and dried. The dry devices were cured overnight at room temperature and stored with the open holes facing downward, in a closed box, until use.

### Sample preparation from staged embryos

Wild-type *C. elegans* embryos (strain N2) were dissected from gravid adults in egg buffer (5 mM HEPES pH 7.4, 118 mM NaCl, 40 mM KCl, 3.4 mM MgCl_2_, 3.4 mM CaCl_2_). Developmental stage was determined by visual inspection of morphology (cell shape and nuclear position) on a dissecting microscope. The embryo with desired stage was transferred to a 3μL drop of lysis buffer (10mM Tris pH 8, 50mM NaCl, 0.1% TX-100, 10% Glycerol), placed the inlet well of a prepared microfluidic device, using a mouth pipet. A clean 26G needle was used to push the embryo into the microfluidic channel.

Once the embryo was trapped in the center of the chamber, the channel output was sealed with crystallography-grade clear tape (Crystal Clear, Hampton Research, Aliso Viejo, CA) to stop flow. The device was temporarily fixed under the dissecting microscope with the tape. The embryo was then immediately crushed while watching in the stereoscope, by pushing down on the surface of the PDMS with the melted tip of a glass Pasteur pipette. A clean glass needle connected to a 10mL syringe through a short flexible tubing was used to puncture the top layer of the PDMS channel once the embryo lysed. The lysate (an approximate volume of 50 nL) was sucked into the needle and transferred onto a marked area of a glow discharged reference grid covered with carbon. Two to three different lysates were transferred onto different squares of the same grid, with no overlap. After the last embryo lysate was transferred, the grid was immediately negatively stained with five consecutive droplets of 2% (w/v) uranyl acetate solution, blotted to remove residual stain, and air-dried in a fume hood.

### Electron microscopy and data collection

Data was acquired using a JEOL 2010F transmission electron microscope operated at 200 keV with a nominal magnification of x60,000 (3.6 Å at the specimen level). Each image was acquired using a 1 s exposure time with a total dose of ~30-35 e^−^Å^−2^ and a defocus between −1 and −2 μm. A total of 1402 micrographs from 7 samples (3 early embryos and 4 late embryos) were manually recorded on a Gatan OneView camera.

7 independent particle stacks were generated from the micrographs of each sample: 341 micrographs of an early-staged embryo sample (E1), 350 micrographs of an early-staged embryo sample (E2), 250 micrographs of an early-staged embryo (E3), 100 micrographs of a late-staged embryo (L1), 111 micrographs of a late-staged embryo (L2), 147 micrographs of a late-staged embryo (L3), and 103 micrographs of a late-staged embryo (L4). FindEM (Roseman, 2004) was used for template-based particle picking with a template selected from reference-free 2D class averages generated from ~10,000 large particles which were manually picked from the E1 dataset. In total, ~81,600 particles were selected from template picking of all datasets. All image pre-processing was done in Appion (Lander et al., 2009). After removing junk particles, 17,070 particles remained for further processing. Particle box size was set to 576 Å x 576 Å. Reference-free 2D class averages were generated with 100 classes using RELION (Scheres, 2012). The 2D class averages of large particles in the embryo lysate were compared with those of purified 40S ribosomes, 60S ribosomes and 26S proteasomes from *Saccharomyces cerevisiae* (a gift from A. Johnson and A. Matouschek) using EMAN. The micrographs of the yeast ribosomes and proteasomes were taken using the TEM procedures above.

For our 40S ribosome reconstruction, 3D classification was performed using RELION to create 10 classes. We used the structure of a purified DNA-dependent protein kinase catalytic subunit as an arbitrary initial model after being low-pass filtered to 60 Å. The top scoring model when compared to the *S. cerevisiae* 40S ribosome structure (EMDB 4214) contained 1466 particles.

For our 60S ribosome reconstruction, a similar strategy was followed. Two independent particle stacks from E1 and E2 were used. The contrast transfer function (CTF) of each micrograph was estimated using CTFFIND4 (Rohou and Grigorieff, 2015). ~37,200 particles were selected by template picking. After removing junk particles, 13,916 particles were left. Particle box size was set to 432 Å x 432 Å. Reference-free 2D class averages were generated with 100 classes. 3D classification was performed to create 8 classes. The structure of a featureless 3D shape with uniform electron density was chosen as an initial model after low-pass filtering to 60 Å. A subsequent round of 3D classification was performed on the same data using a reconstructed 3D class that was most similar to the 60S ribosome as the new initial model. From this classification, the best of three classes was determined by comparison to a *S. cerevisiae* 60S ribosome structure (EMDB 2811) and contained 3,431 particles.

For our 80S ribosome reconstruction, an initial stack of ~9,000 particles in polysome-like structures were manually selected from all datasets combined. After removing junk particles, 5,638 particles remained for subsequent 2D and 3D classification. Particle box size was set to 576 Å x 576 Å. Reference-free 2D class averages were generated with 200 classes. 3D classification was performed to create 2 classes. The top scoring model when compared to a *S. cerevisiae* 80S ribosome structure (EMDB 2858) contained 1,971 particles.

We additionally performed an initial characterization of small particles found in our micrographs. Using a template free Difference of Gaussian particle picker (Voss et al., 2009), ~165,00 particles were selected from datasets E1 and E2. Particle box size was set to 216 Å X 216 Å. After removing junk particles, 126,095 particles were classified using reference-free 2D classification to generate 150 classes.

Data Plotting: to plot the distribution of ribosomes per polysome, we used Violin Plots for plotting multiple distributions (distributionPlot.m) MATLAB function that is publicly available on MathWorks file exchange. The function is available online at: https://www.mathworks.com/matlabcentral/fileexchange/23661-violin-plots-for-plotting-multiple-distributions-distributionplot-m

## Acknowledgements

We thank Kevin Dalby, Arlen Johnson and Andreas Matouschek for purified protein samples; and members of the Dickinson and Taylor labs for helpful discussions. EM data were acquired in the UT Austin Texas Materials Science Institute maintained by K. Jarvis. This work was supported in part by a Welch Foundation Grant F-1938 (to D.W.T.) and NIH R00 GM115964 (D.J.D.). D.W.T and D.J.D. are CPRIT Scholars supported by the Cancer Prevention and Research Institute of Texas (RR160088 and RR170054).

## Supplementary Information

**Figure S1:**
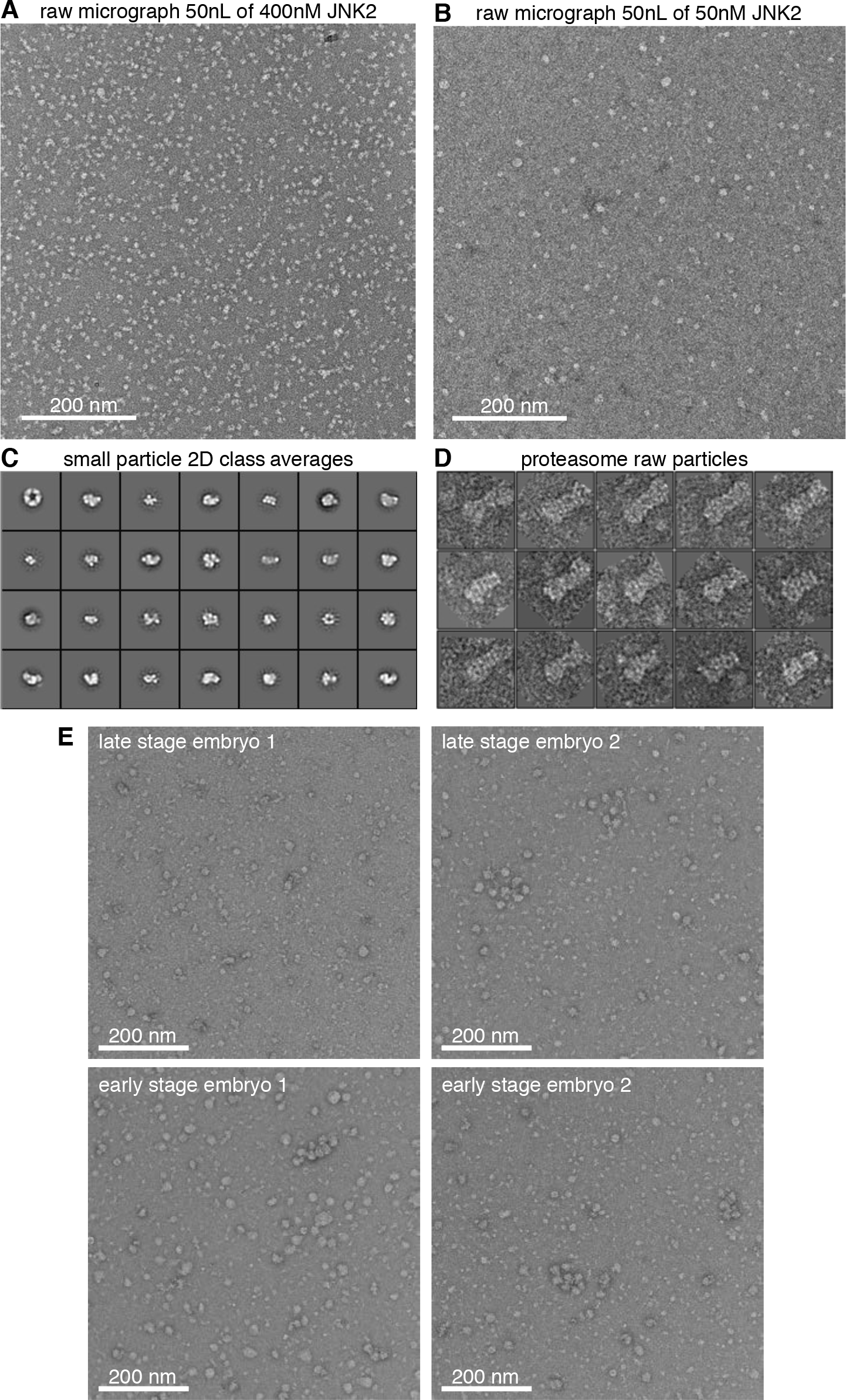
Lysate transfer control experiments and small particle classes. (A)Micrograph of 50nL of 400nM JNK2 (a kinase protein with known structure). To demonstrate that we can visualize particles using our method, we first flowed 400nM JNK2 solution through the PDMS channel and then ~50nL of the solution was transferred to the reference grid using a glass needle. (B)Micrograph of 50nL of 50nM JNK2. (C)Reference-free 2D class averages of small particles picked from two early-staged embryo datasets containing ~126,000 particles. Classes show unique structural features such as a pentameric ring in the top left corner. Individual raw particles of the 26S proteasome show distinct features directly from micrographs. Box size is 576 Å x 576 Å. Representative micrographs from multiple individual single cell experiments shows similar dispersion and size range of single particles.

**Figure S2:**
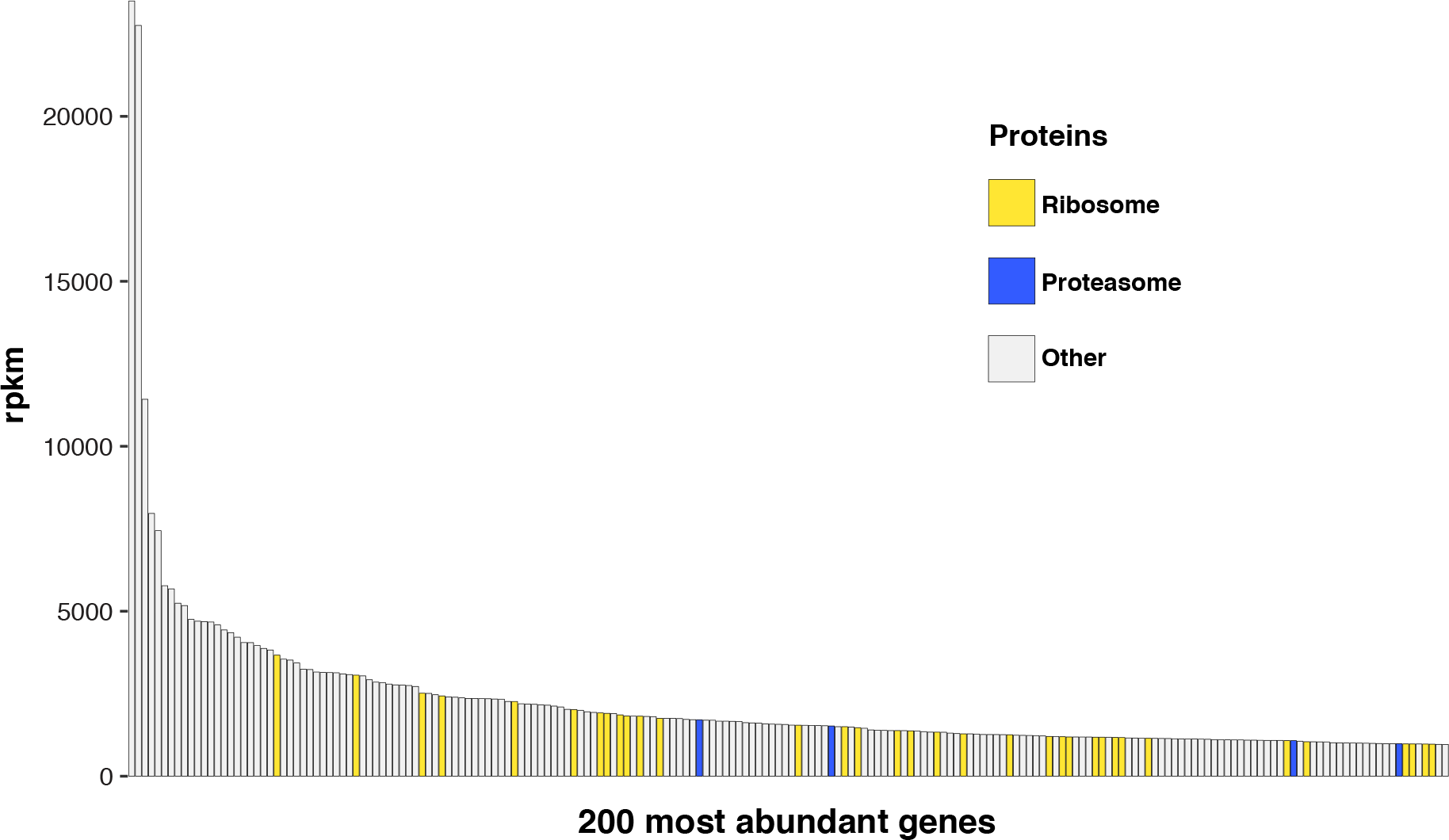
RNA-seq data of *C. elegans* embryos. The 200 most abundant genes in *C. elegans* zygotes (determined by RNAseq; Hillier et al. 2009; Gerstein et al. 2014) sorted in order of decreasing abundance.

**Figure S3:**
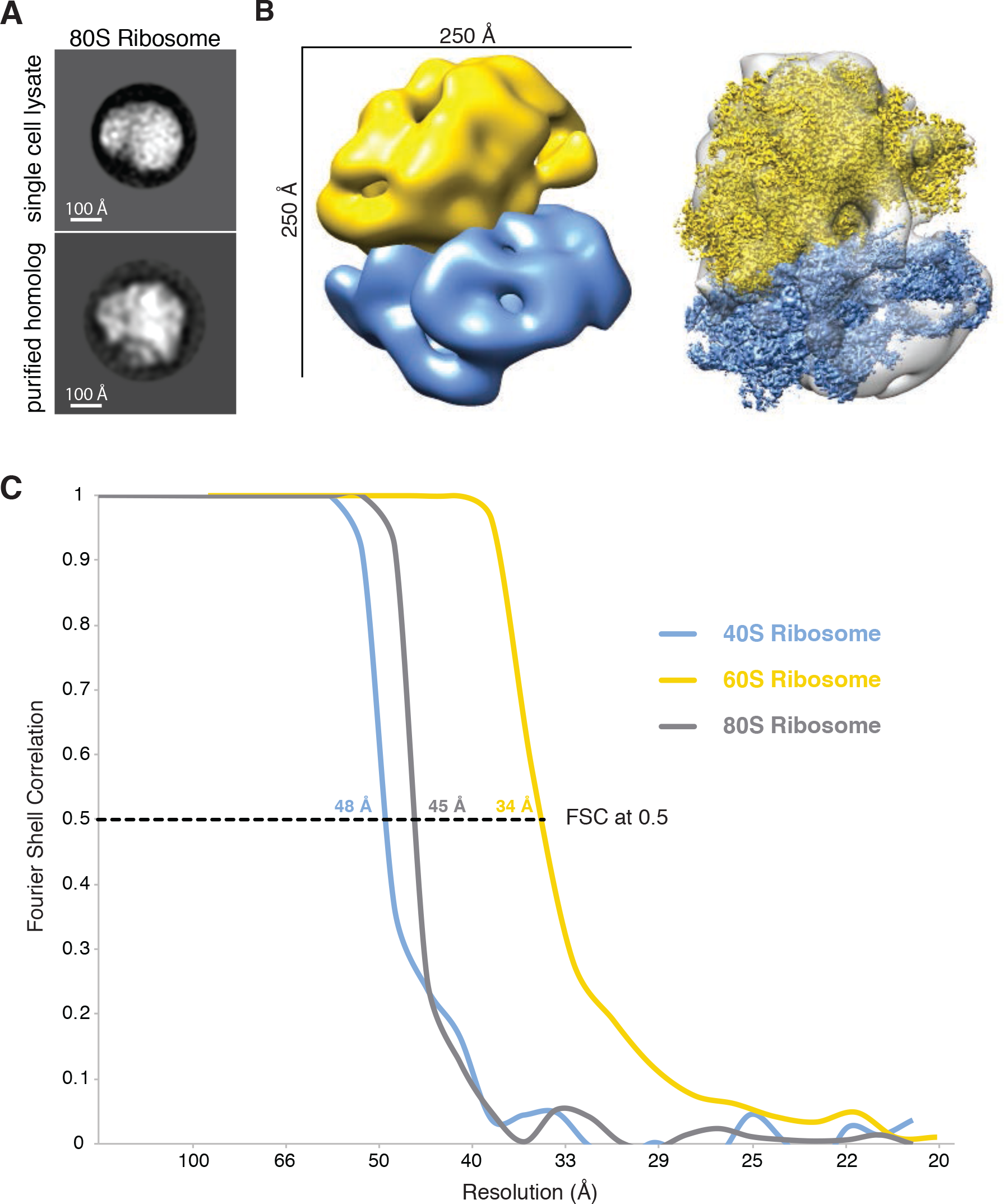
Reconstruction of an 80S ribosome and Fourier shell correlations. (A)Reference-free 2D class average from ribosomes in polysomes aligned to a purified homolog from *S. cerevisiae*. (B)Left: Our 80S ribosome hybrid model. Right: Our 80S ribosome model reconstructed from ribosomes in polysomes with high-resolution 60S (EMDB-2811) (Shen et al., 2015) and 40S (EMDB-4214) (Scaiola et al., 2018) shown in yellow and blue, respectively. (C)Fourier shell correlations of our 40S, 60S and 80S ribosome models. Nominal resolution values are reported at a correlation score of 0.5.

**Supplemental Movie 1**:Video showing the process of embryo lysis and transfer of the lysate to a reference EM grid.

